# A Robust Deep Learning Approach for Joint Nuclei Detection and Cell Classification in Pan-Cancer Histology Images

**DOI:** 10.1101/2023.05.10.540156

**Authors:** Vidushi Walia, Sujatha Kotte, Naveen Sivadasan, Hrishikesh Sharma, Thomas Joseph, Binuja Varma, Geetashree Mukherjee, V.G Saipradeep

## Abstract

Advanced image processing methods have shown promise in computational pathology, including the extraction of crucial microscopic features from histology images. Accurate detection and classification of cell nuclei from whole-slide images (WSI) play a crucial role in capturing the molecular and morphological landscape of the tissue sample. They enable widespread downstream applications, including cancer diagnosis, prognosis, and discovery of novel markers. Robust nuclei detection and classification are challenging due to the high intra-class variability and inter-class similarity of the microscopic morphological features. This is further compounded by the domain shift arising due to the variability in tissue types, staining protocols, and image acquisition. Motivated by the ability of the recent deep learning techniques to learn complex patterns in a bias-free manner, we develop a novel and robust deep learning model TransNuc, based on vision transformers, for simultaneous detection and classification of cell nuclei from H&E stained WSI. We benchmarked TransNuc on the comprehensive Open Pan-cancer Histology Dataset (PanNuke), sampled from over 20,000 WSI, comprising 19 different tissue types and five clinically important cell classes, namely, Neoplastic, Epithelial, Inflammatory, Connective, and Dead cells. TransNuc exhibited superior performance compared to the state-of-theart, including Hover-Net and Micro-Net. TransNuc was able to learn robust feature representations and thereby perform consistently better for the abundant classes such as neoplastic, and the under-represented classes such as dead cells. Similar performance gains were also obtained for epithelial and connective classes that have a significant inter-class morphological similarity.

## I. Introduction

Histopathological images and spatial multiomics play a central role in improved cancer interventions. Integration of histology images, such as H&E stained WSI, and spatial multiomics allows capturing of detailed landscape of the microscopic morphological features and spatial molecular profile of the tumour. Such detailed spatial morphomolecular models allow eliciting the underlying dynamics of the Tumour microenvironment (TME) and thereby enable improved prognosis and discovery of novel markers that aid in personalized interventions. Furthermore, detailed spatio-morphological profiling of WSI can have an immense impact on the diagnostic potential of computational pathology [1]. For instance, accurate detection of tumour and lymphocyte nuclei enables the identification of tumour infiltrating lymphocytes (TILs), which is an important marker for cancer prognosis and recurrence [2]. The nucleus is an object of study not merely to identify cell types, especially in cancer tissues, but also in understanding the cell population, migrating cells, protein localization, phenotype classification, and treatment profiles.

Hematoxylin and Eosin (H&E) staining procedure has been the standard method to distinguish the different tissue types as well as identifying the visible morphological changes in these tissues, as they become cancerous. H&E works well across different fixatives and brings out the various nuclear, cytoplasmic, and extracellular matrix (ECM) features, required for the nucleus detection task. Further, cell and cancer-specific type patterns of heterochromatin condensation can be identified using H&E-stained WSIs [3].

Nucleus detection and cell classification from H&E stained WSI is a highly challenging problem due to multiple factors. There is high intra-class variability and interclass similarity of the microscopic morphological features, including the chromatin patterns. Additionally, there is considerable inter-tumour and intra-tumour heterogeneity of the underlying morphology for the same cell class. Further, the different cell classes appear in the tumour sample with varying degrees of abundances. For instance, the tumour nuclei are abundant in the core and are present in clusters, whereas the dead cells are often sparsely distributed across the tissue. The challenges are further compounded by the domain shift induced by a) variability in staining protocols and image acquisition, b) batch effects, and c) technical artefacts.

Motivated by the potential of machine learning mod-els in computation pathology, several challenges namely, The 2018 Data Science Bowl Challenge [4], CoNSeP[5], MoNuSeg [6], CPM-15, CPM-17[7] and TNBC were organized for building automated nucleus classification and segmentation models. However, these challenge datasets often lacked the diversity that was needed to build robust and generalizable ML models [8]. To address these short-comings, a comprehensive and diverse dataset PanNuke was developed for nucleus detection, classification, and segmentation. The dataset sampled from over 20,000 WSI consisted of 19 different tissue types that were semi-automatically annotated and clinical quality controlled by pathologists. The resulting PanNuke dataset has a high diversity and minimal selection bias, thereby replicating a data distribution that is similar to “the clinical wild” [8]. There are several deep learning and hybrid approaches in the literature for nuclei detection and classification [5], [9]. These approaches can be broadly classified as either, a two-stage approach, where, the nuclei detection or segmentation task is performed first, followed by the classification task, or, a single-stage approach, where both tasks are performed simultaneously. In the following, we discuss the state-of-the-art approaches. The above mentioned approaches mostly rely on CNN based architecture to accomplish the task at hand. However, the introduction of Vision based Transformers have shown improvements over the CNN based architectures by enabling access to more global information [10].

DEtection TRansformer (DETR) [11] is an end-to-end object detection model comprising a convolutional backbone and an encoder-decoder Transformer. DETR has exhibited competitive or superior performance than the state-of-the-art on various (natural) image datasets [11]. A linear layer is added to the DETR architecture for object classification, and an MLP for bounding box prediction. DETR employs a set loss functions based on bipartite matching between the ground truth and the prediction bounding boxes. DETR reduces the dependence on handcrafted modules and post-processing steps namely NMS, making it simple, elegant, and efficient for complex tasks. Our model TransNuc was obtained by training DETR on the PanNuke dataset. To learn feature embedding that is robust to morphological variations and domain shift, we use a variety of image augmentation and normalization techniques. We benchmark TransNuc against state-of-the-art approaches including HoVer-Net [5], Micro-Net [12], Mask R-CNN (MRCNN) [13], Det U-Net [9], DIST [14] and TSFD-Net [15]. Our model was able to learn robust feature representations and thereby perform consistently better than the state-of-the-art for the abundant classes such as neoplastic, and the under-represented classes such as dead cells. Similar performance gains were obtained for epithelial and connective classes with significant inter-class morphological similarity. To the best of our knowledge, our work is the first attempt to establish the potential of vision transformers for both the nucleus detection and classifica-tion tasks on the comprehensive PanNuke dataset.

## II. Materials & Methods

### A. Data

We used the PanNuke [8] dataset which contains 7904 H&E stained image patches sampled from more than 20,000 WSI spanning 19 different tissue types. Nuclei are classified into 5 different clinically important categories, namely, neoplastic cells, inflammatory cells, connective cells, dead cells, and epithelial cells. Nuclei in the dataset have been semi-automatically labelled and further quality-controlled by pathologists. This dataset contains 189,744 labeled nuclei where each nucleus is labeled with an instance segmentation mask. This dataset provides patch-level annotations, with each patch of size 256*×*256 pixels at 40*×* resolution. The original patches, which were scanned at 20*×*, were re-sized to 40*×* for uniformity.

### B. Data pre-processing

Augmentations are applied on training data for robustness and improved generalization. We perform horizontal flipping, vertical flipping, random rescaling, random cropping, and random rotation to make the model invariant to the inter-tissue morphological variations. In [16], the effects of stain augmentation and stain normalization were investigated for robustness and improved generalization performance while processing H&E stained visual fields. Stain augmentation showed improved performance compared to stain normalization [16]. In particular, HSV and HED color transformations, were found to be the key ingredients for improved performance [16]. In [17], using HSV stain augmentation resulted in improved generalization performance for mitosis detection from H&E stained visual fields. In TransNuc, we apply random HSV transformations to randomly change the hue, saturation, and value (HSV) representation of images, to make the model robust to color perturbations caused by variability in image acquisition and staining protocols.

### C. Model Training

The DETR network was trained using the 256×256 patches. Resnet50 backbone pretrained on ImageNet was selected as the first stage, based on ablation experiments. The AdamW optimizer was used for model training. The initial learning rate was set to 1e-04. The batch size was chosen as 6. The hyper-parameters of DETR were chosen using Optuna hyper-parameter optimization framework.

## III. Results

We benchmarked TransNuc with state-of-the-art methods, including, HoVer-Net [5], Micro-Net [12], Mask R-CNN (MRCNN) [13], Det U-Net [9], DIST [14], and TSFD-Net [15]. In the PanNuke dataset, the pre-extracted patches were split into three randomized train, test and validation folds. As recommended by PanNuke, all the models were trained under the same data splits for fair benchmarking. The average of the detection and classification results for the three folds are shown in the Table I. We have also trained TransNuc, where in each fold, the training was done on train and eval splits combined. We denote this model as TransNuc+ in the results Table I.

**TABLE I:**
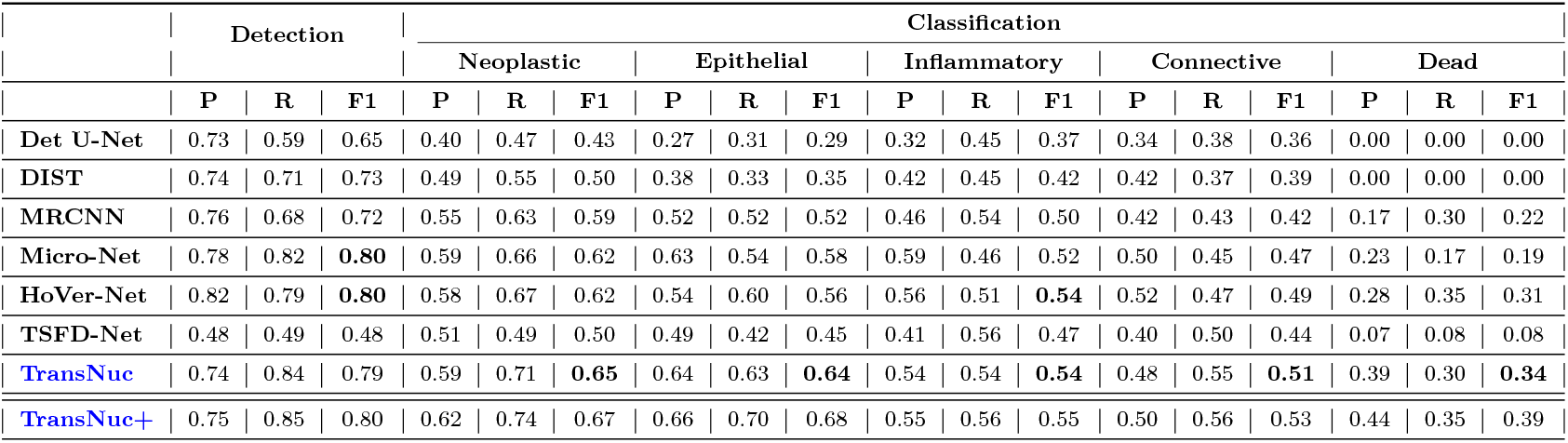
Precision (P), Recall (R) and F1-score (F1) for detection and classification on PanNuke datasets. All results are averaged over the three folds. The highest F1 scores are highlighted. The last row TransNuc+ indicates the performance of TransNuc where, each fold, the training was done on the train and eval splits combined.

|  | Detection |  |  | Classification |  |  |  |  |  |  |  |  |  |  |  |  |  |  |
| --- | --- | --- | --- | --- | --- | --- | --- | --- | --- | --- | --- | --- | --- | --- | --- | --- | --- | --- |
|  |  |  |  | Neoplastic |  |  | Epithelial |  |  | Inflammatory |  |  | Connective |  |  | Dead |  |  |
|  | P | R | F1 | P | R | F1 | P | R | F1 | P | R | F1 | P | R | F1 | P | R | F1 |
| Det U-Net | 0.73 | 0.59 | 0.65 | 0.40 | 0.47 | 0.43 | 0.27 | 0.31 | 0.29 | 0.32 | 0.45 | 0.37 | 0.34 | 0.38 | 0.36 | 0.00 | 0.00 | 0.00 |
| DIST | 0.74 | 0.71 | 0.73 | 0.49 | 0.55 | 0.50 | 0.38 | 0.33 | 0.35 | 0.42 | 0.45 | 0.42 | 0.42 | 0.37 | 0.39 | 0.00 | 0.00 | 0.00 |
| MRCNN | 0.76 | 0.68 | 0.72 | 0.55 | 0.63 | 0.59 | 0.52 | 0.52 | 0.52 | 0.46 | 0.54 | 0.50 | 0.42 | 0.43 | 0.42 | 0.17 | 0.30 | 0.22 |
| Micro-Net | 0.78 | 0.82 | **0.80** | 0.59 | 0.66 | 0.62 | 0.63 | 0.54 | 0.58 | 0.59 | 0.46 | 0.52 | 0.50 | 0.45 | 0.47 | 0.23 | 0.17 | 0.19 |
| HoVer-Net | 0.82 | 0.79 | **0.80** | 0.58 | 0.67 | 0.62 | 0.54 | 0.60 | 0.56 | 0.56 | 0.51 | **0.54** | 0.52 | 0.47 | 0.49 | 0.28 | 0.35 | 0.31 |
| TSFD-Net | 0.48 | 0.49 | 0.48 | 0.51 | 0.49 | 0.50 | 0.49 | 0.42 | 0.45 | 0.41 | 0.56 | 0.47 | 0.40 | 0.50 | 0.44 | 0.07 | 0.08 | 0.08 |
| TransNuc | 0.74 | 0.84 | 0.79 | 0.59 | 0.71 | **0.65** | 0.64 | 0.63 | **0.64** | 0.54 | 0.54 | **0.54** | 0.48 | 0.55 | **0.51** | 0.39 | 0.30 | **0.34** |
| TransNuc+ | 0.75 | 0.85 | 0.80 | 0.62 | 0.74 | 0.67 | 0.66 | 0.70 | 0.68 | 0.55 | 0.56 | 0.55 | 0.50 | 0.56 | 0.53 | 0.44 | 0.35 | 0.39 |

We use the PanNuke specified metrics of Precision, Recall and F1 to compare the detection and classification performances. Specifically, the outputs of all models were evaluated using the PanNuke provided code.^1^ Further details on the metrics are provided in the later sections. A detection is considered to be a true positive if the detection bounding box centroid falls within a 12 pixel radius of the ground truth centroid. The benchmark results are given in Table I. For TransNuc, the centroids of the bounding box predictions were directly supplied to the PanNuke validation script. We have also indicated the performance of TransNuc+ in the results Table I.

Performances of all the competing tools that we have benchmarked, except TSFD-Net, are taken from the Pan-Nuke benchmark study [8], where the tools are evaluated exactly in the manner described above, using the PanNuke supplied data folds and evaluation metrics. Since, TSFD-Net was published post the PanNuke benchmark study [15], for uniform comparison, we trained three TSFD-Net models, one for each of the three data folds. The model checkpoints were based on the best training epochs. The remaining hyperparameters were as recommended by TSFD-Net [15]. For metrics computation, pixel-level segmentation outputs of TSFD-Net were supplied to Pan-Nuke validation scripts, where, the centroids for the predictions are calculated and used for computing the various metrics.

Figure 1 presents a selection of PanNuke image patches with overlayed bounding boxes for both the ground truth and the predictions from the TransNuc. The left side visual fields represent the overlayed ground truth while the right side visual fields show the overlayed predictions from TransNuc. The class of the nuclei is represented by a different color where Neoplastic cells are represented by red color, blue is used for connective cells, orange color represents Epithelial cells, inflammatory cells are marked with green color and yellow color marks the boundaries of dead cells.

**Fig. 1:**
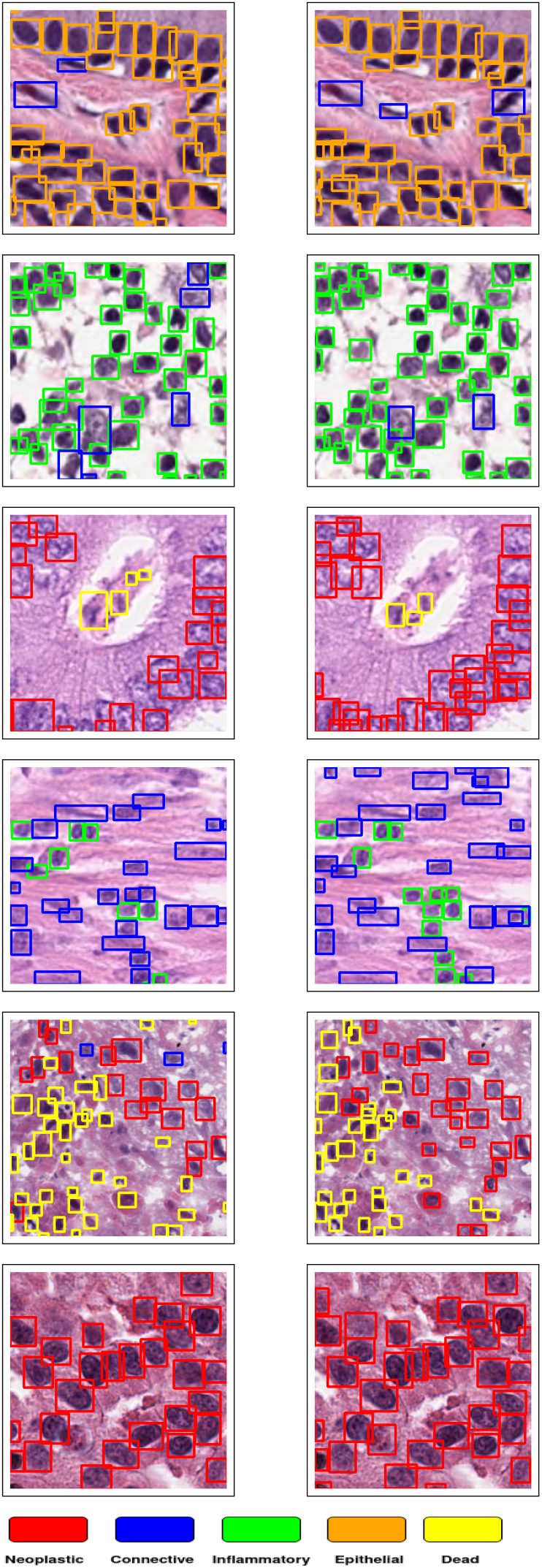
An example of PanNuke visual fields overlayed with ground truth (left column) and TransNuc predictions (right column).

### A. Metrics

For comparing the detection based performance, we adopt the same evaluation metrics as proposed in the PanNuke challenge [5], [8] for nucleus detection and classification. PanNuke benchmark uses precision, recall, and F1-score for evaluating detection and classification performances. PanNuke evaluation method [8], [5] first computes centroids for the predictions and the ground truths. Then, a unique pairing is created between the predicted and the ground truth centroids using Kuhn-Munkres algorithm with distance as the cost function (as proposed by [8], [5]). A detection is paired with a ground truth (a true positive event) if the distance between the prediction centroid and the ground truth centroid is within 12 pixel radius. In [5], [8], the details on computation of precision, recall, and F1 score are provided.

### B. Ablation Studies

1. *Stain Augmentation:* Our experimental results indicate that stain color augmentation improved the object detection performance. Random HSV transformations increased the model’s ability to generalize to unseen stain variations. HSV showed slightly improved performance compared to HED.
2. *CNN Backbones for DETR:* In this study, various experiments were conducted for the choice of DETR backbones (ResNet50, ResNet50-DC5, ResNet101 and ResNet101-DC5). ResNet50 (0.79 F1-score for detection) backbone showed a slight improvement over other back-bones.
3. *Loss functions:* We have used a combination of boundary loss and label loss as our overall loss. Various experiments were conducted on boundary loss functions like DIoU, GIoU, and label loss functions like cross-entropy (CE) and focal loss(FL). Our experiments showed that the model trained with CE loss showed an improvement of 8% in detection performance over the model trained with FL loss function. On the contrary, a change in the boundary loss function from DIoU to GIoU did not improve the detection or classification performance of the model.
4. *Usage of Noisy Priors:* We also experimented with using a noisy prior at the input, where Hematoxylin (H), Eosin (E), and Residual (R) channels from PanNuke images were added as a fourth, fifth, and sixth channels respectively. DETR is trained on these six channel images to check the effect of adding H, E, and R channels. However, our experiments showed adding HER channels did not result in improved detection performance.
5. *Patch size upscaling:* We experimented with training TransNuc after upscaling the input 256 pixel patches to 512 pixels. However, this did not significantly change the detection and classification performance.

## IV. Conclusion

Accurate detection and classification of microscopic cell nuclei from whole-slide images (WSI) have a significant role in capturing the molecular and morphological landscape of the tissue sample, that have applications in several downstream analyses such as cancer diagnosis, prognosis, and discovery of novel markers. The problem is complex due to the high intra-class variability and inter-class similarity of the microscopic morphological features, compounded by domain shift. Our model TransNuc achieved state-of-the-art performance on the comprehensive Pan-Nuke dataset. Our work shows that vision transformers hold promise in fine-grained analysis of WSIs. Future work includes extending our model to handle additional nucleus sub-classes.

## Footnotes

1 https://github.com/vqdang/hovernet/blob/master/computestats.py

